# A Diverse Genetic Basis for Metabolic Reactions is Revealed Through Pangenome analysis

**DOI:** 10.1101/2025.01.23.634541

**Authors:** Omid Ardalani, Patrick V. Phaneuf, Jayanth Krishnan, Gaoyuan Li, Joshua Burrows, Lars K. Nielsen, Bernhard O. Palsson

## Abstract

Sequenced genomes for thousands of strains of a bacterial species allow for a comprehensive analysis of its pangenome. We present a pangenome study of *Escherichia coli*’s metabolism by formulating ‘gene-to-protein-to-reaction associations’ (GPRs) for about 2,700 metabolic reactions in 2,377 fully sequenced strains. On one hand, these GPRs reconstruct strain–specific networks that allow computational predictions (and experimental validation) of metabolic phenotypes, while on the other hand, they give the genetic basis for a given metabolic reaction in every strain. A pangenome-wide analysis of GPRs shows that: 1) We can reveal the genetic basis for a specific metabolic property at the species level; 2) The genetic basis for many metabolic reactions is diverse; 3) Many rare genes show variation in the gene’s genomic neighborhood which often contain genes from transposable elements, 4) Many rare genes show large-scale fragmentation and horizontal gene transfer (>11,000 rare genes in 2,377 strains); and 5) The aromatic amino acids and branched chain amino acids pathways are enriched with rare genes, with Acetolactate synthase having 29 distinct genes. Thus, analysis of GPRs across the pangenome reveals a complex dynamic evolutionary history of metabolism, revealing the role of conserved, fragmented, and horizontally transferred metabolic genes.

## INTRODUCTION

Shortly after the first whole-genome sequences were completed^1–3^, functional annotations enabled the identification of metabolic genes^4,5^. Each gene’s chemical equation could then be used to reconstruct genome-scale metabolic maps tied to underlying genetic data through gene-to-protein-to-reaction (GPR) associations^6,7^, laying the foundation for computational genome-scale models (GEMs) that link metabolic phenotypes to genotypes^8,9,10–12^. GPR associations provide direct links between genotypes and phenotypes, supporting studies on enzyme promiscuity^13^ and flux distribution^14^.

The increased affordability of genome sequencing in the early 2010s led to tens of thousands of bacterial genomes being sequenced, opening new avenues for comparative genomics and pangenomics within bacterial species^15,16^. Pangenomics has since become a powerful addition to phylogenetics, enabling comprehensive analysis of taxonomic genotypes^17,18^. However, despite this wealth of genotypic data, linking genotype to phenotype across pangenomes remains a significant challenge. This challenge can now be addressed at the pangenome scale. We can now formulate panGEMs and panGPRs (pangenome-wide collections of GEMs and GPRs from a large number of sequenced stains of a species). panGPRs expand our ability to compare genotypes and phenotypes across bacterial groups and to infer evolutionary relationships^19–22^.

The *E. coli* pangenome is considered open, with significant genetic diversity across strains expanding as new genomes are sequenced, revealing substantial variability among lineages^23^. The core genome closes at around 2,398 genes, and the accessory genome reaches approximately 5,182 genes, but the number of novel genes steadily increases with new sequenced genomes, driven by genes that are found in single or a handful of strains (called rare genes)^24^. These rare genes provide unique adaptations, allowing *E. coli* strains to thrive in diverse ecological niches^25^.

Frequently acquired through horizontal gene transfer, rare genes enhance metabolic and pathogenic versatility, underscoring their evolutionary importance^26^. Understanding rare genes and their origin is key to decoding the complex evolutionary pathways of *E. coli*. Big data analysis of strain genomes across species thus enables a deeper understanding of evolution and adaptation, as each sequenced strain reflects an outcome of natural evolutionary processes.

## RESULTS

We previously built a pangenome for *E. coli* from a set of 2,337 completely sequenced genomes^27^. The genes in the pangenome can be classified into three categories (see ^28^): 1)the core genome, containing genes found in the vast majority of strains (>96.7%), 2) the accessory genome, containing genes found in 6.8% to 96.7% of strains, and 3) the rare genome that contains genes found in less than 6.8% of strains. As with the genes, metabolic reactions can be classified similarly into core reactions, accessory reactions, and rare reactions. Thus, relationships between metabolic genes and reactions can be studied at the pangenome scale using GPRs.

**Figure 1.**
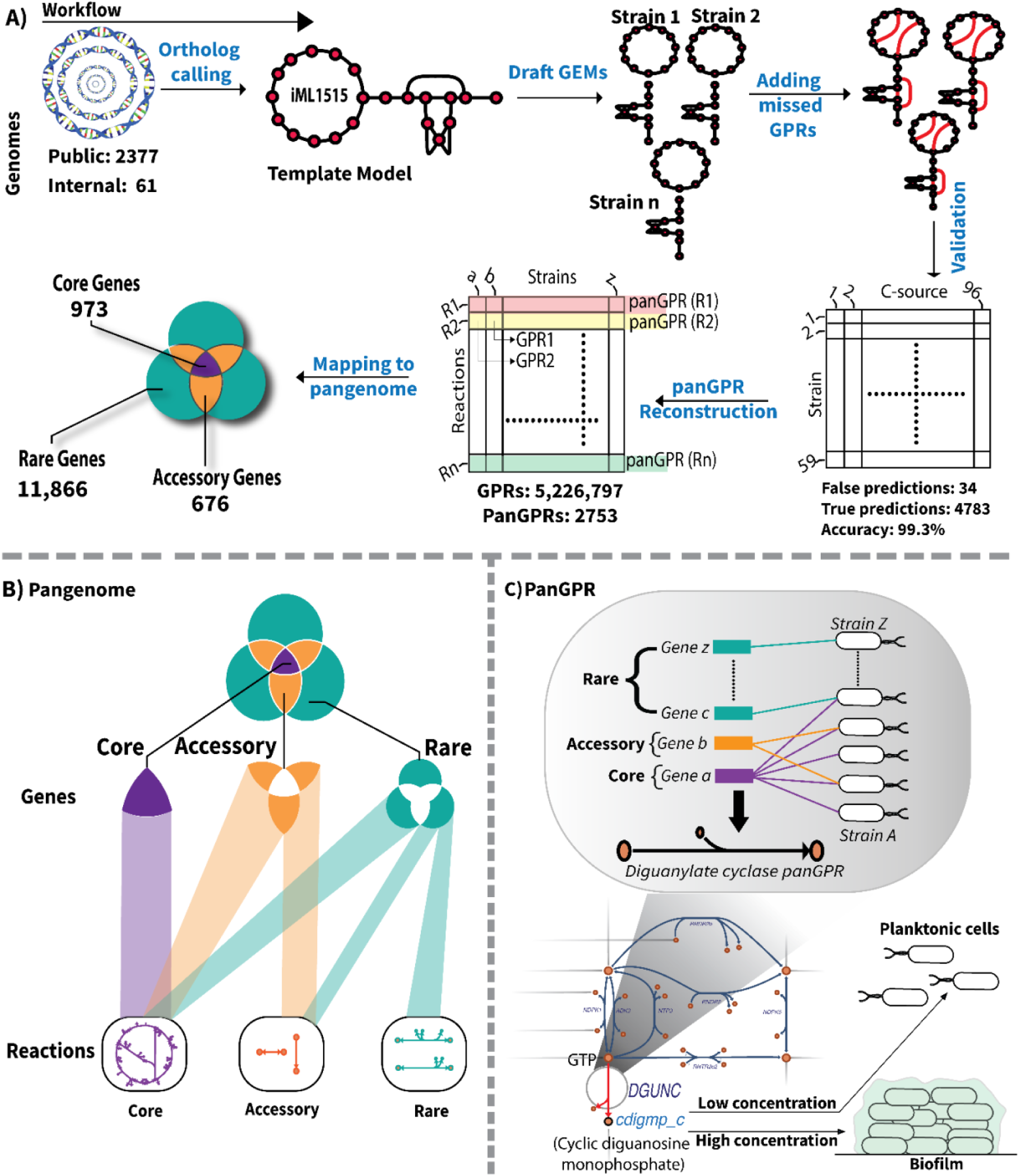
Generating the panGPRs. **A)** Workflow. A total of 2,377 high-quality E. coli genomes were annotated using PROKKA and mapped to the iML1515 metabolic reconstruction available on the BiGG database to generate strain-specific draft networks. These draft networks were then curated semi-automatically, followed by a manual curation step that included curating reactions and metabolite namespaces and bidirectional BLAST against all GEMs of Enterobacteriaceae members available on the BiGG database, followed by adding missing reactions from the KEGG database to generate a curated panGEM. The panGEM was validated by generating GEMs for a subset of 59 in-house strains whose genomes were assembled in contigs and comparing their growth phenotypes on 96 carbon sources with BioLog growth phenotypes (Figure 2, below). The curated and validated panGEM was then mapped to an existing pangenome of E. coli to formulate the panGPRs. **B)** Overview of how the PanGPRs originate from the pangenome. Analyzing panGPRs revealed that 94% of rare metabolic genes, which are found in only a few strains, encode core metabolic reactions that are found in all GEMs of the panGEM. **C)** PanGPRs for the case of diguanylate cyclase. The panGPRs capture all possible genetic make-ups that encode each metabolic reaction. Analyzing these genetic make-ups revealed that the genetic basis of reactions is diverse due to the overlapping functions of rare and accessory genes with core genes, with reactions involved in cell surface modulation and biofilm formation having the most diverse genetic basis. The case of Diguanylate cyclase (DGUNC) that regulates the switch between plantonik and biofilm life styles is shown.

### panGPRs define a pan-genetic basis for E. coli metabolic reactions

The gold standard set of GPRs from *E. coli* K-12 MG1655^12^ was mapped onto the 2,337 complete genomes to generate 5,226,797 GPRs (see Methods). This set of GPRs formed the first version of the panGPRs for the *E. coli* species. Next, all PROKKA annotations identifying metabolic genes on the 2,377 genomes, to which no MG1655 GPR was associated, were mapped. Then, using bi-directional BLAST (see Methods) to match all GPRs found in the BiGG Models database (<u>bigg.ucsd.edu</u>), we found an additional 7,412 metabolic GPRs for 7,720 genes from other species found in this database formulated into eight distinct reactions. For 67,054 genes, we constructed 35,549 additional GPRs (formulated into 30 distinct reactions) based on the KEGG database (<u>genome.jp/kegg/pathway.html</u>). This yielded the set of 5,269,758 GPRs for the metabolic genes in the pangenome, which together constitute the panGPRs used in this study (see list in SI File 1).

### A panGEM links metabolic genotypes to phenotypes with a high degree of accuracy

The completeness of the GPRs for a strain is assessed through metabolic network reconstruction followed by the formulation of computational genome-scale models (GEMs) of metabolism^9^. A panGEM was built by mapping the panGPRs onto the 2,377 completely sequenced genomes to reconstruct strain-specific metabolic networks. This panGEM contains 2,753 distinct metabolic reactions, which included 2,025 metabolites and 13,515 distinct metabolic genes, comprising 973 core, 676 accessory, and 11,866 rare metabolic genes.

In addition, GEMs were built in the same fashion for a set of 59 fully sequenced and phylogroup-balanced in-house *E. coli* strains. These 59 GEMs were used to predict metabolic states that can be compared to measured metabolic phenotypes (Fig 2A). The metabolic phenotypes were measured using BioLog plates where 11,712 growth conditions were generated in replicates (16,997=59 strain *96 wells *3 replicates).

**Figure 2.**
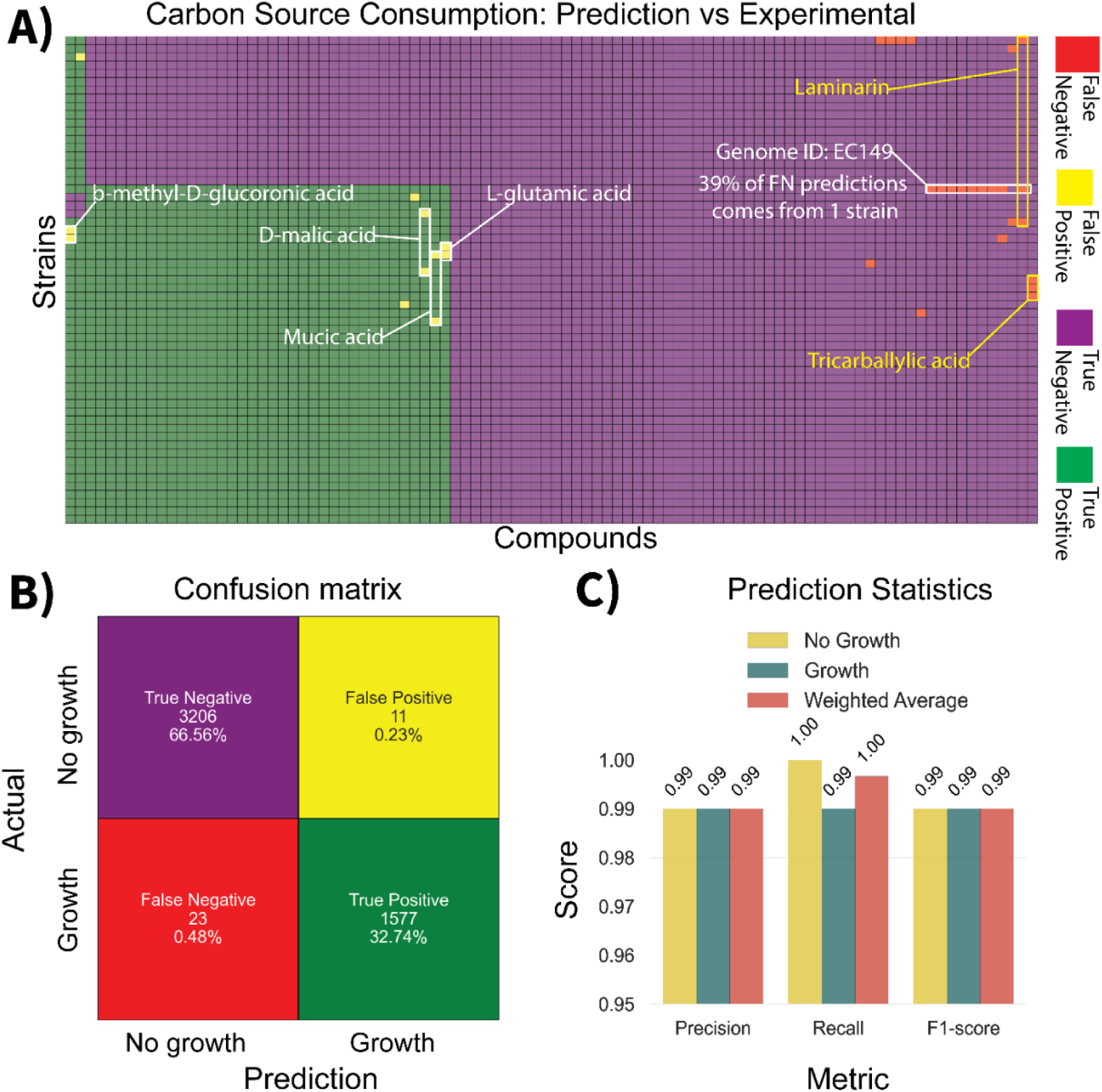
Validation of panGEM predictions. **A)** Comparison of Predicted and Experimental Carbon Source Consumption: The X-axis represents 96 compounds, while the Y-axis displays 59 strains (see Extended Data Fig 1 for a detailed description of columns and rows). Each cell in the heatmap is color-coded according to the provided legend, illustrating discrepancies or alignments between predicted and experimental data. Note that most of the false negatives are associated with one strain (strain ID EC149). **B)** Carbon Source Consumption Confusion Matrix: This matrix details the counts and percentages for True Negatives, False Negatives, True Positives, and False Positives, providing a quantitative overview of prediction accuracy. **C)** Prediction Statistics: Bar plot displays the precision, recall, and F1-scores derived from the validation results, offering a statistical summary of the predictive performance. See detailed description in main text.

The confusion matrix (Fig 2.B) provides a detailed breakdown of the panGEM’s prediction performance. True negatives (3206 instances, 66.56%) are correctly identified as ‘no growth’ conditions, while true positives (1,577 instances, 32.74%) are correctly identified as ‘growth’ conditions. False positives (11 instances, 0.23%), predict growth where none is observed and false negatives (23 instances, 0.48%), predict no growth where growth is observed. The combined results indicate a low error rate in the panGEM’s predictions.

Additional prediction statistics further underscore the panGEM’s robustness, with precision, recall, and F1-score metrics all averaging around 0.99 (Fig 2.C). This result suggests that the panGEM not only accurately predicts ‘growth’ and ‘no growth’ conditions but does so consistently across different strains. The high precision indicates that the panGEM’s predictions for growth are mostly correct, while the high recall ensures that the majority of actual growth conditions are identified. The F1-score, which balances precision and recall, confirms the overall effectiveness of the panGEM.

This large-scale evaluation demonstrates panGEM’s predictive capabilities. This high accuracy indicates that the quality of i) the genomic sequences, ii) their annotation, iii) the assembled panGPRs, and iv) the metabolic network reconstruction is high, leading to reliable predictions.

### Phylogroup distribution of metabolic genes and reactions in the panGEM reveals conserved metabolism across phylogroups

A workflow using these predicted phenotypes classifies the GEMs for the 2,377 strains into phylogroups (Extended Data Fig. 2). GEMs exhibit extensive variability in metabolic gene counts across different phylogroups, reflecting genetic diversity and adaptability. Shigella strains, a named phylogroup of *E. coli*, have lower metabolic gene counts compared to other *E. coli* phylogroups. This result is consistent with their genome reduction resulting from an adaptation strategy of obligate pathogens that leads to the loss of unnecessary genes^29^ (see Extended Data Fig. 2 and Extended Data Fig. 3 for details.).

While the median metabolic gene count is relatively similar across phylogroups, notable outliers are found in all phylogroups (Extended Data Fig. 2). As an example, phylogroup E, including 285 strains, stands out with 70 rare reactions, indicating less conserved metabolism. Phylogroup Shigella, including 49 strains, has 44 rare reactions, also suggesting a less conserved metabolic profile. In contrast, phylogroups C and B1, which have the highest count of core reactions among all phylogroups, exhibit the most conserved metabolism (Table S1).

Although no metabolic reaction was found to be specific to a phylogroup, their genetic basis was phylogroup-specific (Extended Data Fig. 4). The phylogroup-specific reactomes (a set of reactions that can be found across strain-specific GEMs of a phylogroup) are similar in their metabolic reaction content, but the genetic basis for their reactions can be diverse. In other words, the phylogroup-specific reactomes represent a similar set of reactions, but the genes they are associated with may differ, as detailed below.

### Defining phylons: groups of genes that co-segrate in the pangenome

Phylons have recently been defined based on non-negative matrix factorization (NMF) of the pangenome matrix^27^. The rows in the pangenome matrix represent all the genes found in the pangenome, and the columns represent the individual sequenced genomes. An entry of ‘1’ is placed into this matrix if the corresponding gene is found in the corresponding genome. Otherwise, the entry is ‘0’. The pangenome matrix is thus a binary matrix, succinctly representing the gene portfolio of all the strains used for the analysis.

NMF finds groups of genes that are found together in a set of strains and absent in other strains, thus defining the ‘phylons.’ Phylons therefore modularize the pangenome into sets of genes that are found as a group in a strain but absent in other strains. A total of 31 phylons are found in the *E. coli* pangenome^27^. They are consistent with previous phylogroup designations and subsets thereof. Phylons thus represent the genetic basis for phylogroups. The metabolic genes specific to a phylon are of interest here.

### Phylon-based clustering of GEMs reveals the genetic diversity underlying conserved metabolic functions

The genetic basis for metabolic reactions across the phylons exhibits considerable diversity in gene content while simultaneously highlighting the conservation of core metabolic functions (Fig 3). The panGEM was analyzed to define the location of core, accessory, and rare metabolic reactions across the phylons. The metabolic reaction corresponding to these three classes of genes was mapped to the *E. coli* pangenome. The results from this mapping reveal that the majority of metabolic genes belong to the rare category (11,866 genes). In contrast, most metabolic reactions that are historically classified as belonging to core metabolic functions (2,711 core reactions) are associated with genes in the core genome. Surprisingly, panGPR analysis indicated that 94% of rare metabolic genes also encode reactions in core metabolic pathways (Fig 3A, inset).

**Figure 3.**
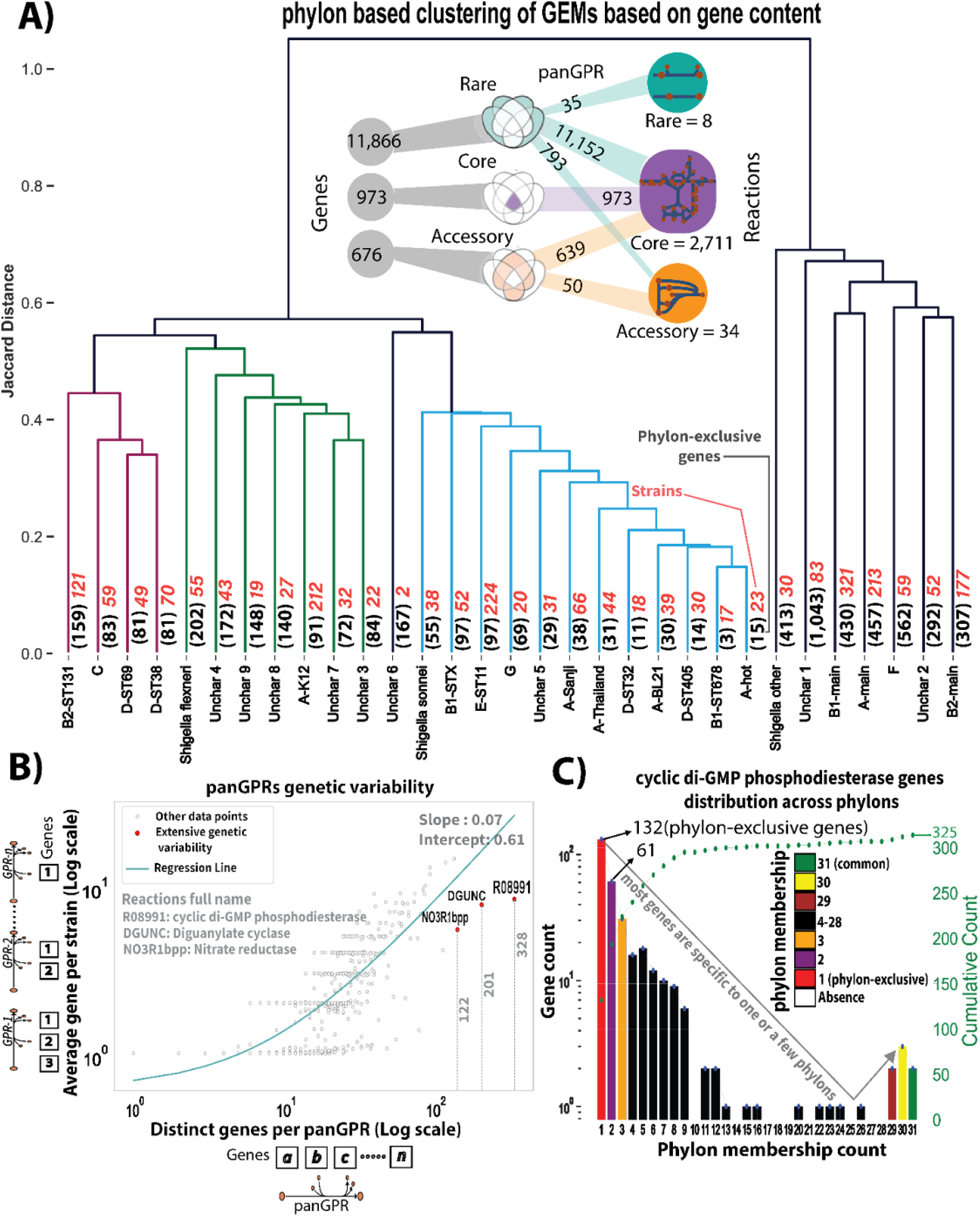
Phylon-specific Gene to Protein to Reaction (GPR) Associations for metabolic genes. **A)** Hierarchical clustering of metabolic genes across the phylons. The phylons are named after the most similar phylogroup (i.e., C, G, F), sequence type (ST) within a phylogroup (i.e., B2-ST131, etc.), or are ‘uncharacterized’ (these pylons often represent mobile elements, see ^27^). Each branch terminates with a set of phylon-specific metabolic genes, with unique gene counts indicated in parentheses. The numerical values in red represent the number of strains constructed for each phylon. The inset graphic displays the distribution of reactions and genes (using the panGPRs), which are categorized as core, rare, or accessory, highlighting their count within the dataset and the proportion of gene categories in reaction categories. **B)** The scatter plot demonstrates the relationship between the count of distinct genes per reaction (i.e., in all GPRs for a reaction) and the average gene count per GPR. The x-axis represents the number of distinct genes coding for enzymes catalyzing the same reaction, while the y-axis indicates the average number of genes per reaction within GPRs of each reaction. This plot highlights reactions with high diversity in their genetic basis such as, cyclic di-GMP phosphodiesterase (R08991), diguanylate cyclase (DGUNC), and nitrate reductase (quinone-dependent) (NO3R1bpp). **C)** The histogram illustrates the distribution of genes across phylons, with cumulative counts representing the total genetic variants for R08991 reaction. The y-axis shows the number of distinct genes encoding R08991, while the x-axis displays the number of phylons in which each gene is found. The bars are color-coded to indicate gene frequency.

Based on these results we can assess the presence of phylon-specific metabolic reactions. No metabolic reactions were found to be exclusive to a single phylon. However, the analysis identified unique metabolic genes specific to each phylon (Extended Data Fig. 4). Notably, the ‘Uncharacterized 1,’ ‘F,’ and ‘A-main’ phylons were found to harbor the highest numbers of unique metabolic genes, with 1,043, 562, and 457 genes, respectively. This finding suggests that despite the observed genetic diversity of metabolic genes, core metabolic reactions and functions remain conserved across different phylons (Fig 3A). This result shows that the diversity of metabolic genes is greater than that of the corresponding metabolic reactions.

### The pangenomic basis for some metabolic reactions exhibits extensive genetic variation

A deeper examination of the genetic basis underlying each metabolic reaction revealed significant variability in the genes encoding these functions across the panGEM (Fig 3B) (full list in SI File 2). We observed that these genes are enriched in functions related to cell surface modulation, which play a key role in the transition between planktonic and biofilm states. Among the reactions with the most variable genetic basis are cyclic di-GMP phosphodiesterase (R08991) (Fig 3B) (see Extended Data Fig. 5 for protein similarity analysis), diguanylate cyclase (DGUNC), and quinone-dependent nitrate reductase (NO3R1bpp). They exhibited the highest genetic diversity, with 328, 201, and 122 distinct genes, respectively, across the panGEM. Intriguingly, the majority of the genes for these three metabolic reactions were phylon-specific (Extended Data Fig. 6) or are found in the rare genome. This result underscores the genetic heterogeneity associated with specific metabolic functions related to biofilm formation. Cyclic di-GMP is a well-known second messenger in prokaryotes. Elevated cytosolic levels of cyclic di-GMP signal biofilm formation. The reactions DGUNC and R08991 synthesize and degrade cyclic di-GMP, respectively, modulating the transition between planktonic and biofilm life style. Additionally, NO3R1bpp can use nitrate as a terminal electron acceptor under low oxygen conditions during biofilm formation^30–33^.

### Many strains have rare genes associated with the tryptophan biosynthesis pathway

The reactions in the tryptophan biosynthesis pathway show an enrichment of rare genes, including two critical reactions representing commitment steps at branch points, both of which exhibit high genetic variability. Rare genes were identified for all three isozymes of 3-deoxy-D-arabino-heptulosonate 7-phosphate synthetase (DDPA), the first commitment step in the aromatic amino acid (AAA) biosynthesis pathway. Specifically, we found 11 rare genes encoding *aroH* across 41 strains, 9 rare genes encoding *aroG* across 19 strains, and 10 rare genes encoding *aroF* across 25 strains. While *aroG* and *aroF* each have one core gene and no accessory genes, *aroH* lacks a core gene but includes three accessory genes, suggesting that a core function can be maintained without the presence of a core gene. Instead, different strains may encode the same enzymatic function using rare and accessory genes.

Anthranilate synthase (ANS), which catalyzes the first commitment step in tryptophan biosynthesis, shows a high prevalence of rare genes. ANS is a heterotetrameric complex composed of two *trpD* and two *trpE* subunits. Both subunits exhibit a notable count of rare genes, with *trpE* encoded by 8 rare genes across 205 strains, and *trpD* by 7 rare genes across 35 strains. Interestingly, *trpE* lacks a core gene but has one accessory gene, while *trpD* contains one core gene and no accessory genes. Similar genetic diversity is found in genes that encode enzymes in other aromatic amino acid biosynthetic pathways as well as those in branched-chain amino acid (BCAA) synthesis (see Extended Data Fig. 7). We note that these amino acids are commonly auxotrophic ^34^.

### Contrasting genomic neighborhoods: stability in core and accessory genes, diversity in rare genes

In a previously published study^27^, we determined that about 80% of genes in the rare genome were transposable elements (TE), enriched in 320 TE families. Some of these TE families carried passenger genes^35^. The remaining 20% of rare genes were of diverse origins. To establish evidence of horizontal gene transfer (HGT) among rare metabolic genes, we analyzed the genomic neighborhood of core, accessory, and rare metabolic genes for all metabolic reactions found in the panGPRs. The results revealed that rare metabolic genes have markedly more variable genomic neighborhoods as compared to core and accessory genes (Figure 4A). The genomic neighbors of core and accessory genes were found to be highly conserved, with one new neighbor gene found per 100 to 1,000 strains. In contrast, rare genes exhibited far greater variability, with one new neighbor gene observed every 1 to 10 strains. This indicates that rare genes’ neighbors vary significantly from strain to strain, while core and accessory genes tend to maintain the same neighbors across hundreds or even thousands of strains.

**Figure 4.**
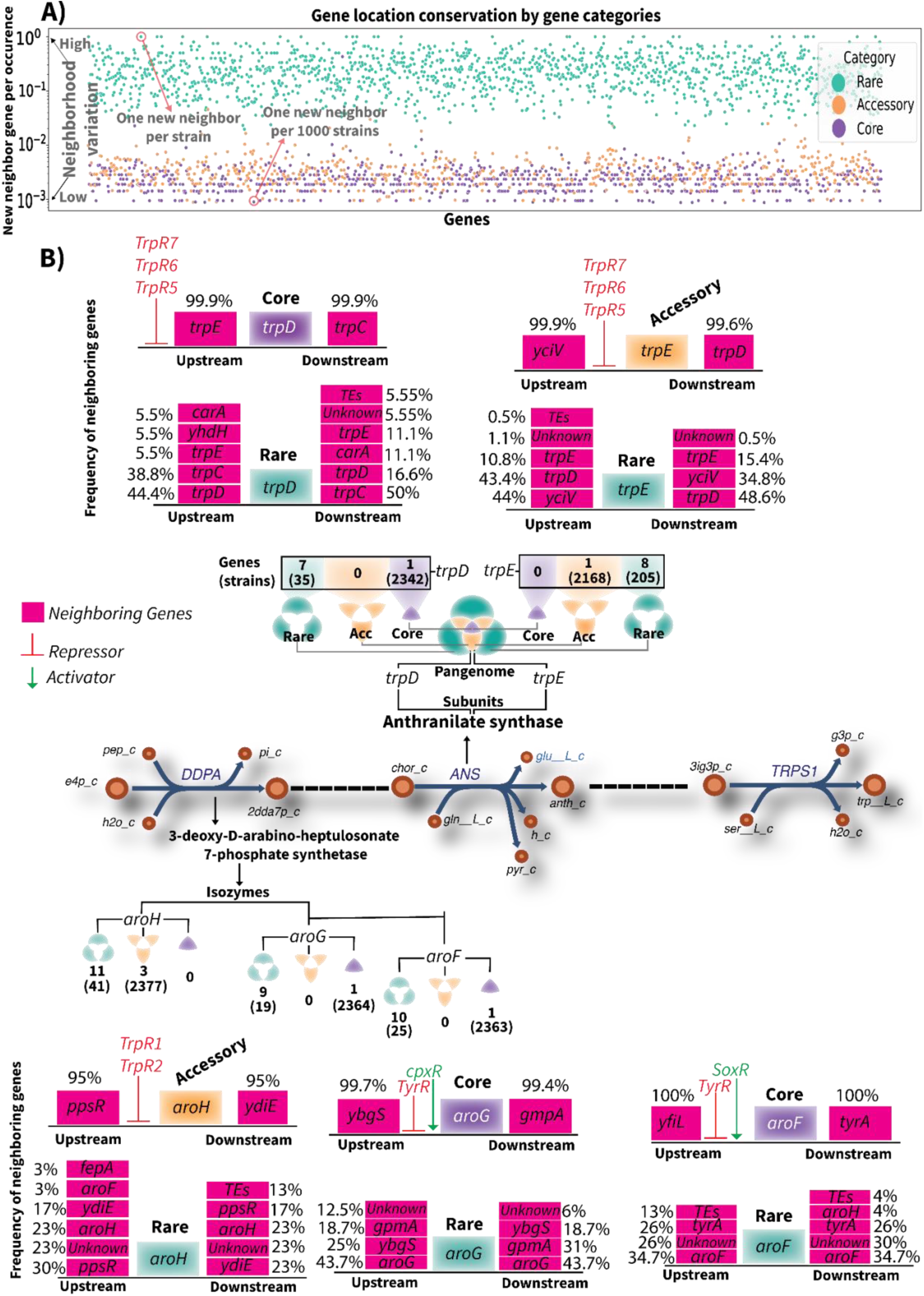
Genome neighborhood for core, accessory, and rare genes with a detailing of this information for the tryptophan biosynthesis pathway. **A)** The dot plot represents the conservation of neighboring genes for core, accessory, and rare genes across all panGPRs. The X-axis shows the genes, and the Y-axis represents the count of distinct neighboring genes per occurrence of the target genes (i.e., close to 1 is highly variable, the lower the y-value the more conserved the genome neighborhood is). Genes are color-coded as core, accessory, and rare genes). **B)** Rare genes involved in the aromatic amino acid biosynthesis pathway. The two commitment steps to aromatic amino acid biosynthesis (DDPA reaction) and then to tryptophan biosynthesis (ANS reaction) are highlighted for a detailed presentation of their genetic basis. Genetic variation in each reaction’s subunits or isozymes are depicted by the number of genes classified as core, rare, or accessory, with the values in parentheses indicating the number of strains in which each category of gene is found. For each isozyme or subunit, the gene’s neighborhood is shown separately for core, rare, and accessory genes to emphasize the genomic location variability of rare genes.

We studied the specific genome neighborhoods of the rare genes found in the tryptophan biosynthetic pathway in more detail. The gene neighborhoods for DDPA isozymes are depicted in Figure 4B. The results show that for all three accessory genes of *aroH*, 95% of strains have conserved neighbors, whereas rare genes display a wide range of neighborhoods, including associations with genes found in transposable elements, unknown proteins, gene duplications, and integration into different transcriptional units. In contrast, the core genes of *aroG* and *aroF* show a 99% to 100% conservation of their neighbors. Rare genes, like those of *aroH*, demonstrate significant neighborhood diversity. A comparable pattern was observed for the subunits of ANS (Figure 4B).

### Various protein families underlie genetic diversity of rare metabolic genes and their phylogenetic origin

A multiple sequence alignment (MSA) was conducted to compare rare genes with core and accessory genes sharing the same functional annotations. The results indicate that the majority of rare genes are likely products of core or accessory gene fragmentation, often located near transposable elements. A smaller subset of rare genes, however, exhibited substantial sequence divergence compared to core and accessory genes. Notably, these divergent genes were found outside the native transcriptional units of their core and accessory counterparts, suggesting horizontal gene transfer (HGT) events (see Extended Data Fig. 8 for detailed analysis of cyclic di-GMP phosphodiesterase genes). For example, among the 11 rare genes encoding aroH in the DDPA panGPR, 10 were found to be fragments of three accessory genes, while one gene (Ecoli_C41084) showed significant sequence divergence (Fig. 5A) and was dislocated from its original transcriptional unit (Fig. 5B) and its genomic region (Fig. 5C). Analysis of the genomic neighborhood of aroH revealed the presence of insA6, insB6, insC6, insD6, insE5, insF5, and IS630 family transposases near the fragmented rare genes, highlighting their potential role in gene fragmentation (Fig. 5B). Similar patterns were observed for aroF, aroG (other DDPA panGPR isozymes) (Extended Data Fig. 9), and the ANS panGPR subunits trpD and trpE, as shown in the Extended Data (Extended Data Fig. 10).

**Figure 5.**
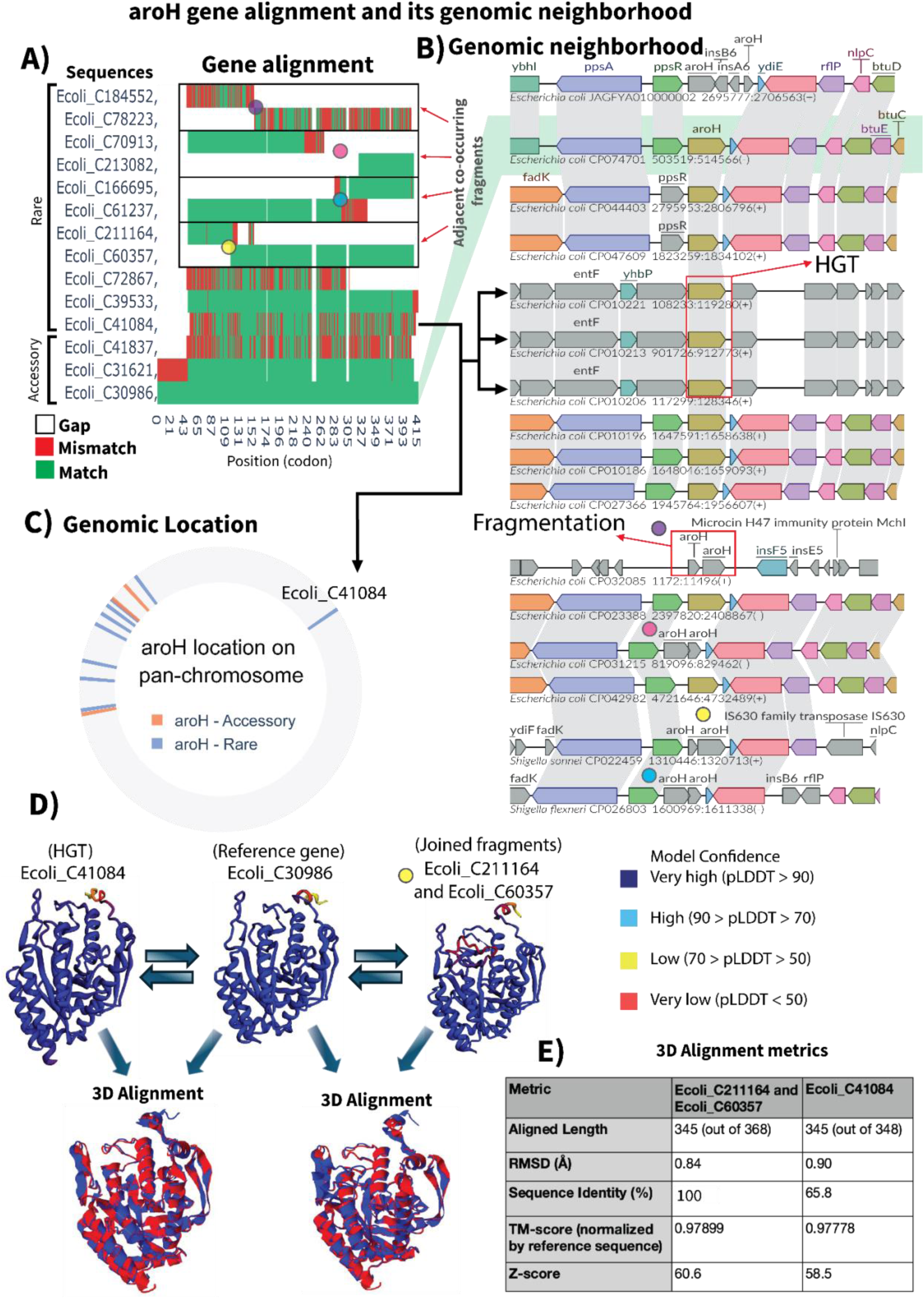
aroH gene alignment and its genomic neighborhood. **A)** MSA of rare aroH genes to accessory aroH gene. Alignment is across the 415 codons in the gene. The C numbers are names of gene clusters, see ^46^. Green represents the codon sequence in the reference (last row). Green segments for other gene clusters indicate consistency with reference sequence, red segments indicate deviation from the reference sequence, and white means these codons are missing. Genes grouped together within a black box represent fragments that are adjacent on the genome (neighboring genes). Each pair of neighboring fragments is marked with colored dots to indicate their positions on the genome, which are highlighted later in panel (B). **B)** genomic neighborhood of accessory and rare aroH genes (corresponding to panel A as shown). Gray shading connects homologous genes across the genome neighborhoods. The anchor gene, AroH, is shown in the brown color). The reference genome neighborhood has a green background. Many of the rare genes have a similar genomic neighborhood as the reference, but some are highly different, as those pointed by a black arrow. The colored dots in panel (B) correspond to the neighboring fragments previously displayed in panel (A). **C)** Genome location of accessory genes are highlighted in orange and rare genes are in blue. Genomes are aligned by the genes shared by all the genomes and only appear once. **D)** 3D protein structure modeling and alignment: The 3D models of protein structures were generated for the reference gene (Ecoli_C30986), a rare horizontally transferred gene (Ecoli_C41084), and the combined fragments of two rare neighboring genes (Ecoli_C211164 and Ecoli_C60357). The 3D alignment compares the rare gene structures against the reference gene. Structures are color-coded based on model confidence. In the 3D alignments, the structure of the reference gene is shown in blue, while the structure of the rare gene is shown in red. **E)** presents the metrics for the 3D alignment of the rare genes against the reference gene. RMSD: Root Mean Square Deviation. pLDDT: per-residue measure of local confidence. TM-score: template modeling score.

To assess structural and functional similarities and differences between the reference aroH gene and rare variants, functional 3D model structures were predicted using the RoseTTAFold server. Model confidence scores were 0.85 for Ecoli_C211164 and Ecoli_C60357 (combined fragments), 0.91 for the reference gene Ecoli_C30986, and 0.89 for the rare horizontally transferred gene (Ecoli_C41084). These predicted structures were subsequently used for 3D alignment (Figure 5D).

The structural alignment of the combined rare fragments (Ecoli_C211164 and Ecoli_C60357) against the reference gene (Ecoli_C30986) revealed complete sequence identity (100%) across aligned regions, (Figure 5E). The TM-score, normalized by the reference sequence, was 0.9879, indicating highly conserved structural features, with an RMSD of 0.68 Å, reflecting minimal divergence in atomic positions.

Similarly, the alignment between the rare horizontally transferred gene (Ecoli_C41084) and the reference gene (Ecoli_C30986) demonstrated remarkable structural similarity. Despite differences in genomic context and sequence, the TM-score was 0.9978, nearly identical to the reference, with an RMSD of 0.50 Å. These results underscore the conserved structural properties of aroH-encoded proteins, even in fragmented or horizontally transferred gene contexts (Figure 5D and E).

## DISCUSSION

We present a pangenome study of *Escherichia coli*’s metabolism. Affordable sequencing has generated a large number of strain-specific sequences. Each sequence represents the outcome of a natural evolutionary ‘experiment.’ We can now collect all these outcomes and perform pangenomic analysis. In addition, prior knowledge gained through metabolic reconstruction allows us to identify the genes that are associated with metabolic reactions across the pangenome of a species through a collection of GPRs. We refer to this pangenome-wide collection of GPRs as the panGPRs for the set of sequenced genomes used in the study. A combination of the panGPRs and the collection of whole-genome sequences thus allows us to perform, for the first time, a detailed species-level analysis of the genetic basis of *E. coli* metabolism.

The analysis reveals a surprising genetic diversity underlying a subset of common metabolic reactions. Although many common reactions have a relatively uniform genetic basis, reflected in genes in the core genome, we find that a fraction of genes in the rare genome encode enzymes catalyzing reactions in common metabolic pathways. This variation is reflected in enzymes of different protein families, but with the same catalytic function, and in a variable genome neighborhood of the gene encoding the enzyme. Thus, this observed genetic diversity does not result in different metabolic pathways.

Our analysis thus reveals surprising genetic diversity underlying conserved metabolic functions. This diversity is indicative of complex evolutionary dynamics of gene loss and gene re-acquisition to maintain the same biochemical function needed for the survival of a strain specialized to one environmental niche when it moves to another. Perhaps one reason for the diversity of the genetic basis of metabolic reactions is the need to rebuild the loss of metabolic pathways that were eliminated as auxotrophy developed. Such strains would not survive the release from a specialized microenvironmental niche unless they re-acquired genes that complete the corresponding biosynthetic pathway.

Our analysis of the aromatic and branched-chain amino acids supports such a ‘gene loss-reacquisition’ mechanism. The tryptophan biosynthetic pathway exhibits considerable genetic variation in its enzymes across the pangenome (Figure 4). We found that the same is true for phenylalanine and tyrosine, the other two aromatic amino acids. We also found that the same is true for the three branched-chain amino acids (BCAA), valine, leucine, and isoleucine. We note that auxotrophies are common amongst the AAAs and the BCAAs as a result of niche adaptation^34^. This result indicates that these auxotrophies are the results of gene fragmentation. When auxotrophic strains leave this niche, they need to re-acquire missing genes in these pathways through horizontal gene transfer.

Horizontal gene transfer can be examined by studying a gene’s genomic neighborhood. We find that the genomic neighborhood of rare metabolic genes is highly variable and in some cases contains genes from transposable elements. Such genomic locations, being quite different from the genome neighborhood of conserved genes catalyzing the same reaction, are strongly indicative of horizontal gene transfer events. While the majority of rare metabolic genes arise from the fragmentation of core or accessory genes, a small subset of rare genes shows significant differences from core or accessory genes, suggesting the possibility of HGT events. Assessment of the essentiality of these rare genes would be additional support for a gene loss/re-acquisition mechanism.

The *E. coli* panGPR represents a notable advance in our understanding of the genetic basis for individual metabolic reactions in this iconic model organism. The unprecedented scale and depth of our analysis have uncovered novel features of the *E. coli* pangenome. Our findings of extensive horizontal gene transfer and gene fragmentation events related to the rare genome (while the genomic location of core and rare genes remains relatively constant) reveal fundamental characteristics of the dynamic nature of bacterial evolution.

## METHODS

### Genome selection criteria

Public genomes: A set of 2773 high quality genomes was retrieved from our previous pangenome study of E. coli^36^.

### Genome Annotation

All downloaded genomes were re-annotated using PROKKA v1.14.5^37^ for consistency in gene annotation when generating the PanGEM.

### Pangenome generation

We used the generated pangenome from our recent study ^36^ which was generated for the same set of genomes as this paper. Briefly, to construct the pangenome, genomes were clustered based on Mash distances, filtering out the top 1% with the highest Mash distance for each group. Hierarchical clustering was then applied with a 0.1 threshold, resulting in 31 clusters that informed NMF decomposition. Core, accessory, and rare genes were defined using a cumulative gene distribution plot, identifying the core at the “elbow” point (90% from inflection to endpoint) and rare genes similarly from the lowest endpoint.

### PanGPRs reconstruction

PanGPRs were generated by mapping selected genomes to iML1515 GPRs from the BiGG database^38^. First, the iML1515 model was downloaded from BiGG models. Second, all GPRs were collected from the model, and discrepancies in reaction IDs and metabolite IDs were corrected to prevent errors in passing them to panGPRs. GPRs were then aggregated to merge genes associated with similar reactions into a single reaction containing all gene variants, referred to as panGPRs. The final set of all panGPRs was called the panreactome. To retrieve the genetic basis of GPRs, locus tags for all mentioned GPRs were collected from the GEMs using cobrapy^39^ and were mapped to their corresponding genomes to obtain their sequences. Obtained gene sequences and panreactome were later used as templates for orthologs calling during panGEM reconstruction.

### PanGEM reconstruction

The panreactome was used as a template for pangenome scale reconstruction of strain-specific GEMs. Panreactome gene sequences and biochemical information were fed into our previously generated pipeline^19^ alongside re-annotated target genomes, and the similarity threshold for ortholog calling was set to 90% for mapping genes of target genomes to GPRs gene sequences using bidirectional BLAST. The output GEMs were then used to get a list of all locus tags in each model. These locus tags were then mapped against their respective genomes to find missed metabolic genes from their corresponding models. Missing genes were re-mapped to BiGG and KEGG databases using their annotation for finding their metabolic information. Genes with existing reactions in their corresponding models were added to their corresponding GPRs, while genes with novel reactions underwent manual formulation of their reactions according to available protocol^10^ and were added to models. The biomass objective function was obtained from IML1515 model^12^ and was added to all GEMs. GEMs then passed a gap-filling procedure on M9 media using our previous method ^19^. After gap-filling, flux balance analysis was performed on M9 media to confirm the capability of generated models in producing biomass.

### Validation

To calculate the precision and accuracy of PanGEM, we selected 59 strains from our in-house collection and performed BioLog phenotype microarray screening on a variety of carbon sources with M9 as base media. All experiments were performed in triplicate. Inconsistent growth/no growth on each carbon source for each strain was filtered out to obtain a set of high-confidence validation data. Results were compared to computational growth prediction under the same simulated condition. *In silico* growth/no-growth results were defined with a threshold of 0.001 h-1 growth rate as a no-growth phenotype. A confusion matrix for the enumeration of false, true, positive, and negative was formed, and accuracy and precision were prepared and shown in Figure 3.

### Biolog PM data processing

For each well in the PM^TM^ dataset, we applied a Savitzky-Golay filter to each signal to smooth them and computed the maximum respiration value observed, the maximum respiration rate, the time taken to reach the observed maximum respiration, value and the area under the curve. For each plate with a negative control well, we defined a threshold for rejecting samples with a strong positive control signal. Based on these filtered samples, we then obtained a distribution of maximum respiration values recorded in the control wells. This control distribution was used to run a z-test/t-test against each subsequent well in every plate to determine if the observed signal was significant enough to be considered a metabolically active well. For plates in which the control well signals were determined to be significant, we manually went through the samples to check if these were cases of control well contamination or a case of high background noise based on strain characteristics. In the case of the latter, we re-analyzed the samples by performing a t-test between wells and control wells on the same plate and reincorporated them into the final dataset. Finally, to determine if a strain can utilize a substrate in a well, we looked at the consensus of yes/no calls on the substrate across all replicates. If the majority of replicates exhibit a positive signal on the substrate, a 1 (yes) is assigned, otherwise 0 (no). A 0.5 (uncertainty) is assigned in cases where the replicates do not show a consensus. An interactive interface to parse and view this phenotype microarray dataset along with the genomes used for model validation is available at https://pmkbase.com/species?specie=ecoli

### Media simulation and computational growth prediction

for the simulation of M9 media, formulation and exchange bounds were retrieved from a previously published study^40^, maximized biomass production was set as the objective function of models, and FBA was performed to predict the growth rate of models on different carbon sources.

### Mapping GPRs to the pangenome

Genes of PanGEM GPRs were mapped to our previously reconstructed pangenome^36^ by their locus tags to identify the gene cluster of each locus tag. Subsequently, all gene clusters coding for the same reaction were identified and grouped based on their reactions to reconstruct panGPR, which depicts the genetic basis of each reaction across the pangenome of *E. coli*.

### Gene genomic neighborhood analysis

The genomic location of each gene was extracted from GenBank files using Biopython to narrow down the visualization to the location of interest. For each gene of interest, genomes were selected to ensure that core, rare, and accessory variants of the target gene were represented in the chosen subset. LoVis4u Version: 0.0.10^41^ was used as the visualization tool to visualize the genomic neighborhood of target genes.

### Gene pan-chromosome location analysis

We first extracted strain vectors from the genome GFF files, with each vector comprising a list of all genes within a strain’s chromosome, ordered according to their genomic positions (i.e., start and end coordinates). To align gene orders across strains, we identified single-copy core genes (present exactly once and shared by all strains) to serve as reference points for alignment. Next, we aligned all the genomes to construct a pan-chromosome. The distance between each single-copy core gene pair on the pan-chromosome was generated by the average number of genes between them across all genomes. The location of each gene was determined by the average distance to the single-copy core gene ahead of this gene.

### Multiple sequence alignment

For the comparison of rare, accessory, and core genes, we performed MSA using Clustal Omega^42^ and Biopython^43^. In cases where core genes were available, they were used as the reference sequence to which rare and accessory genes were aligned. When core genes were absent, an accessory gene was selected as the reference. Gene sequences were obtained in FASTA format, with each gene’s alleles organized under their corresponding gene ID and locus tag. Sequences were aligned using Clustal Omega, and the longest representative sequence for each gene was selected for alignment.

### 3D Structure Prediction and Alignment

The 3D protein structures were predicted using the RoseTTAFold server ^44^, which generates high-confidence models based on sequence information and structural homology. Model predictions were performed for the reference gene (Ecoli_C30986), the rare horizontally transferred gene (Ecoli_C41084), and the combined fragments of two rare neighboring genes (Ecoli_C211164 and Ecoli_C60357). The confidence scores for the predicted structures were 0.91 for Ecoli_C30986, 0.89 for Ecoli_C41084, and 0.85 for Ecoli_C211164 and Ecoli_C60357. The predicted structures were visualized to ensure quality and used for downstream structural alignment.

Structural alignments were conducted using US-align (Universal Structural alignment)^45^. The alignments compared the combined rare fragments (Ecoli_C211164 and Ecoli_C60357) and the rare horizontally transferred gene (Ecoli_C41084) against the reference gene (Ecoli_C30986). Alignment metrics, including sequence identity, TM-score, RMSD (Root Mean Square Deviation), and aligned length, were calculated to quantify structural similarity. TM-scores were normalized using the reference structure to ensure consistency. The results were visualized and summarized in tabular format to highlight structural conservation across rare and reference gene contexts.

## Code and data availability

Generated GEMs are available on Zenodo:

GEMs : https://zenodo.org/records/13825392

Codes and data : https://zenodo.org/records/14028473

Codes are available at github repository EcoPanGEM: https://github.com/omidard/EcopanGEM

## Acknowledgment

We thank Marc Abrams and Daniel Zielinski for reviewing the manuscript and providing suggestions.We thank Siddharth Chauhan for the discussion on E. coli pangenome. This work was supported by grants from the Novo Nordisk Foundation (NNF20CC0035580).

## Author Contributions

Bernhard O. Palsson, Lars K. Nielsen, and Omid Ardalani conceptualized and designed the study. Patrick V. Phaneuf and Omid Ardalani set up high-performance cloud computation. Omid Ardalani reconstructed, curated and validated the panGEM and panGPRs. Jayanth Krishnan conducted the BioLog phenotype microarray screening and processed the data. Gaoyuan Li analyzed genomic locations on the pan-chromosome. Joshua Burrows and Omid Ardalani performed multiple sequence alignments, and Omid Ardalani conducted genomic neighborhood analysis and performed the 3D structure modelling and alignment. All authors contributed to editing the manuscript and approved the final version.

## Ethics declaration

The authors declare no competing interests.

## Extended Data Figures

**Extended Data Fig 1.**
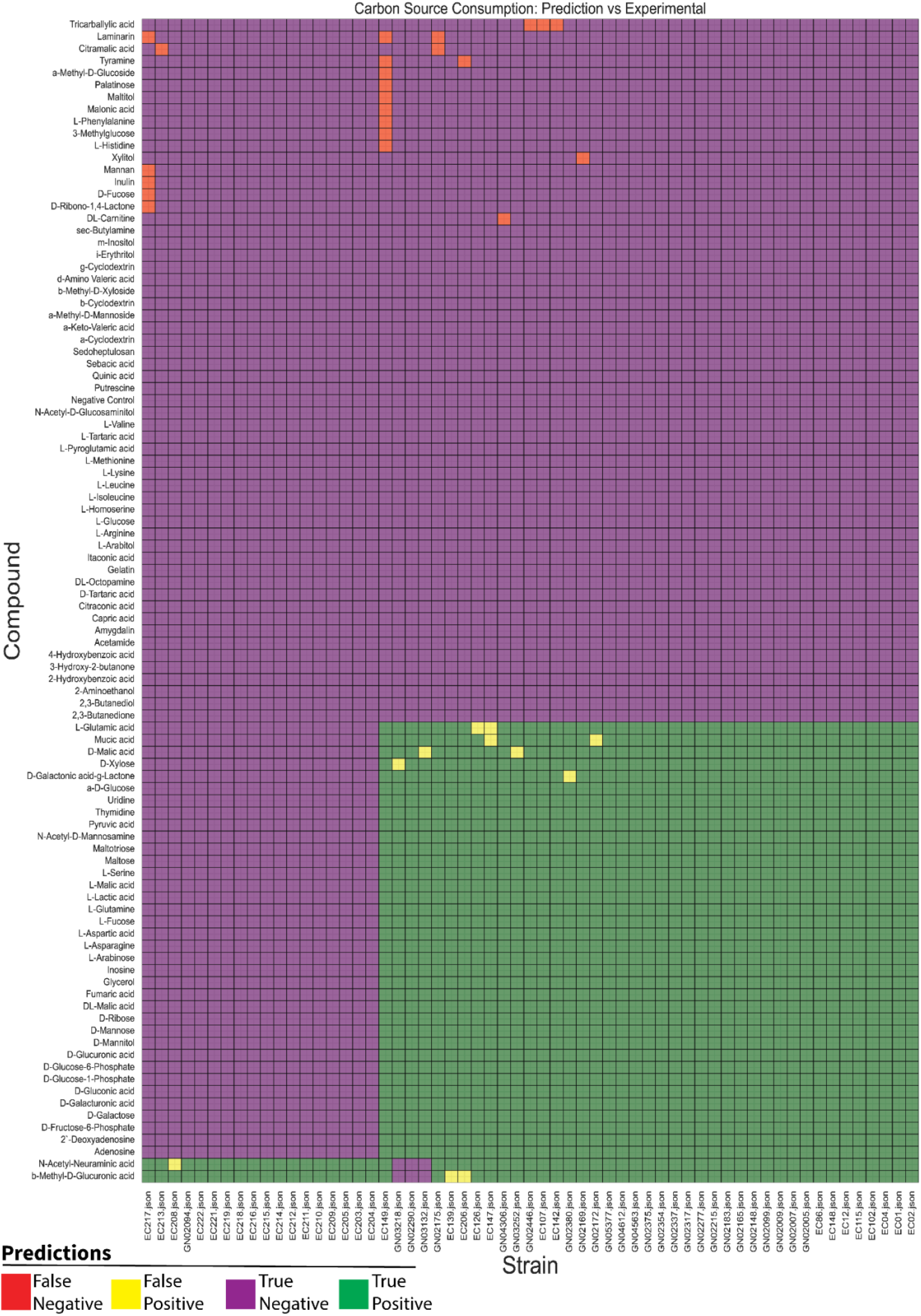
GEM validation. This figure is an enlarged version of Figure 2A to improve readability. The growth of 59 strains was experimentally measured on 96 carbon sources, and the results were compared to the model’s predictions for the same set of carbon sources. Green cells represent true positive predictions, where the model correctly predicts growth that matches the experimental phenotype. Purple cells represent true negative predictions, where the model correctly predicts no growth, consistent with the experimental results. Yellow cells represent false positives, where the model predicts growth, but the strain does not grow experimentally. Red cells represent false negatives, where the model predicts no growth, but the strain grows experimentally.

**Extended Data Fig 2.**
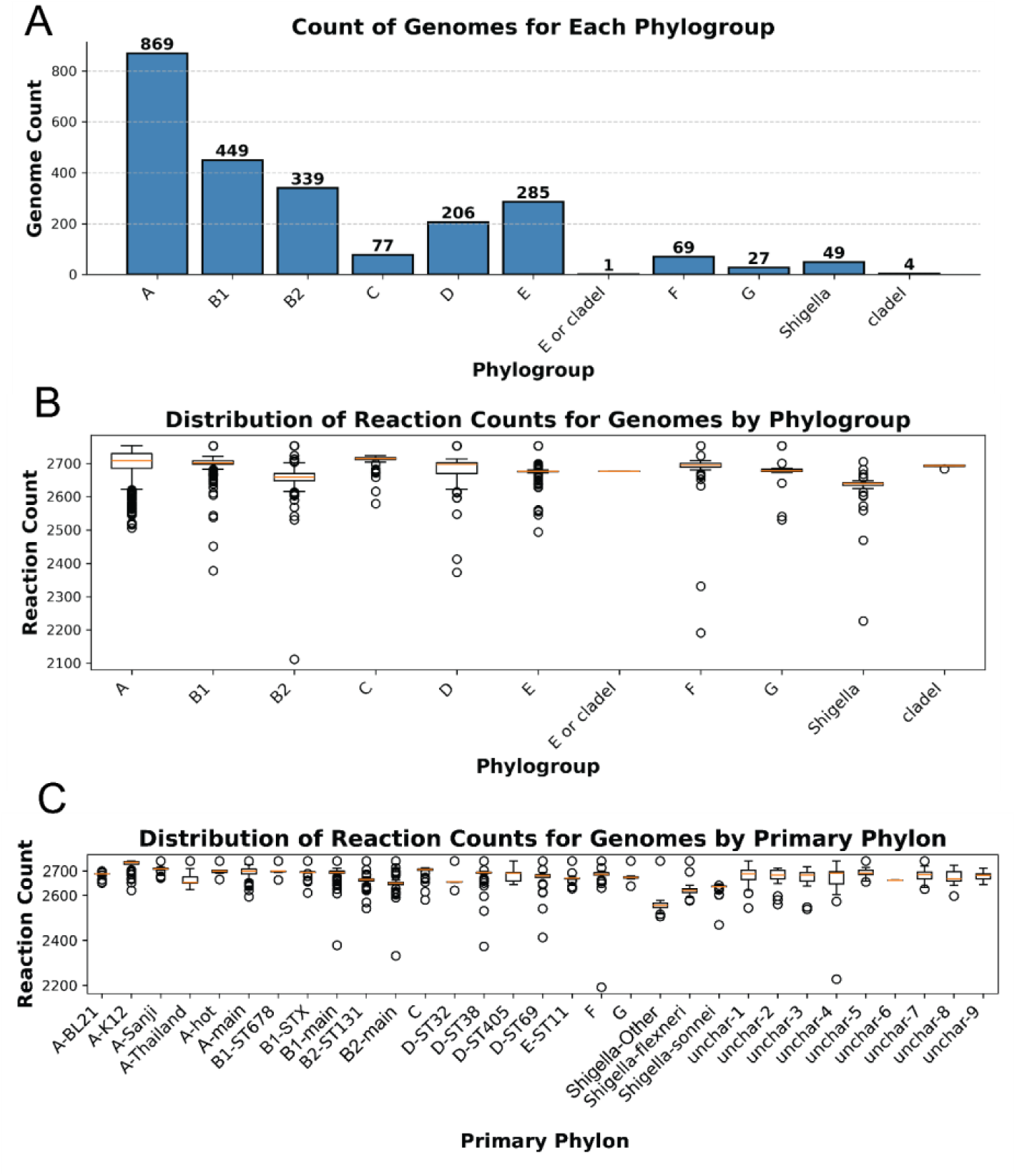
Basic GEM information. A) A bar chart showing the distribution of genomes for which GEMs were reconstructed across different phylogroups. B) A boxplot illustrating the average and range of reaction counts for GEMs within each phylogroup. C) A boxplot illustrating the average and range of reaction counts for GEMs within each primary phylon.

**Extended Data Fig 3.**
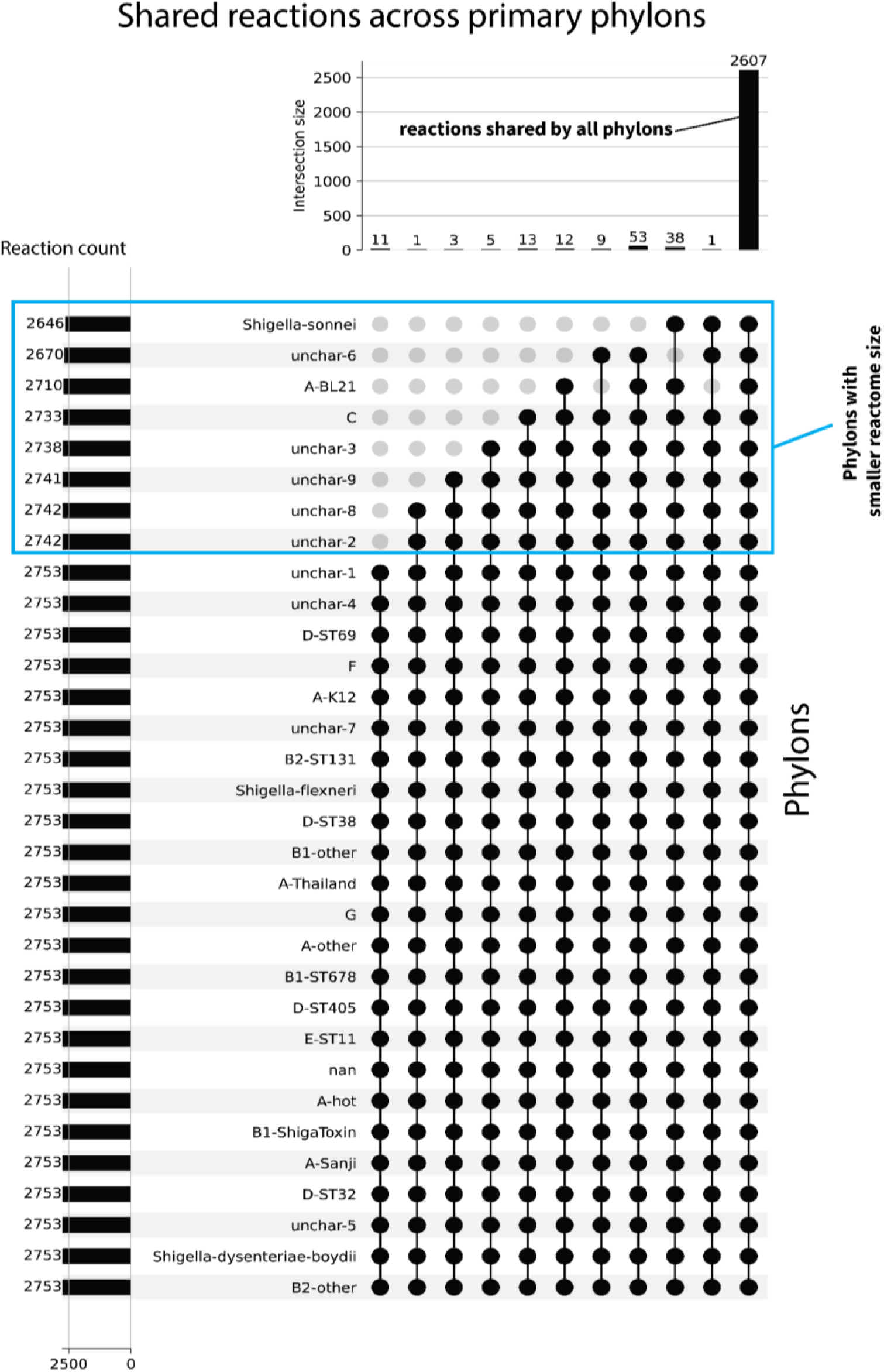
Primary phylons’ reactome size and shared reactions across phylons. The UpSet plot illustrates the intersection of reactions across different primary phylons, showing the number of shared reactions among members of each phylon.

**Extended Data Fig 4.**
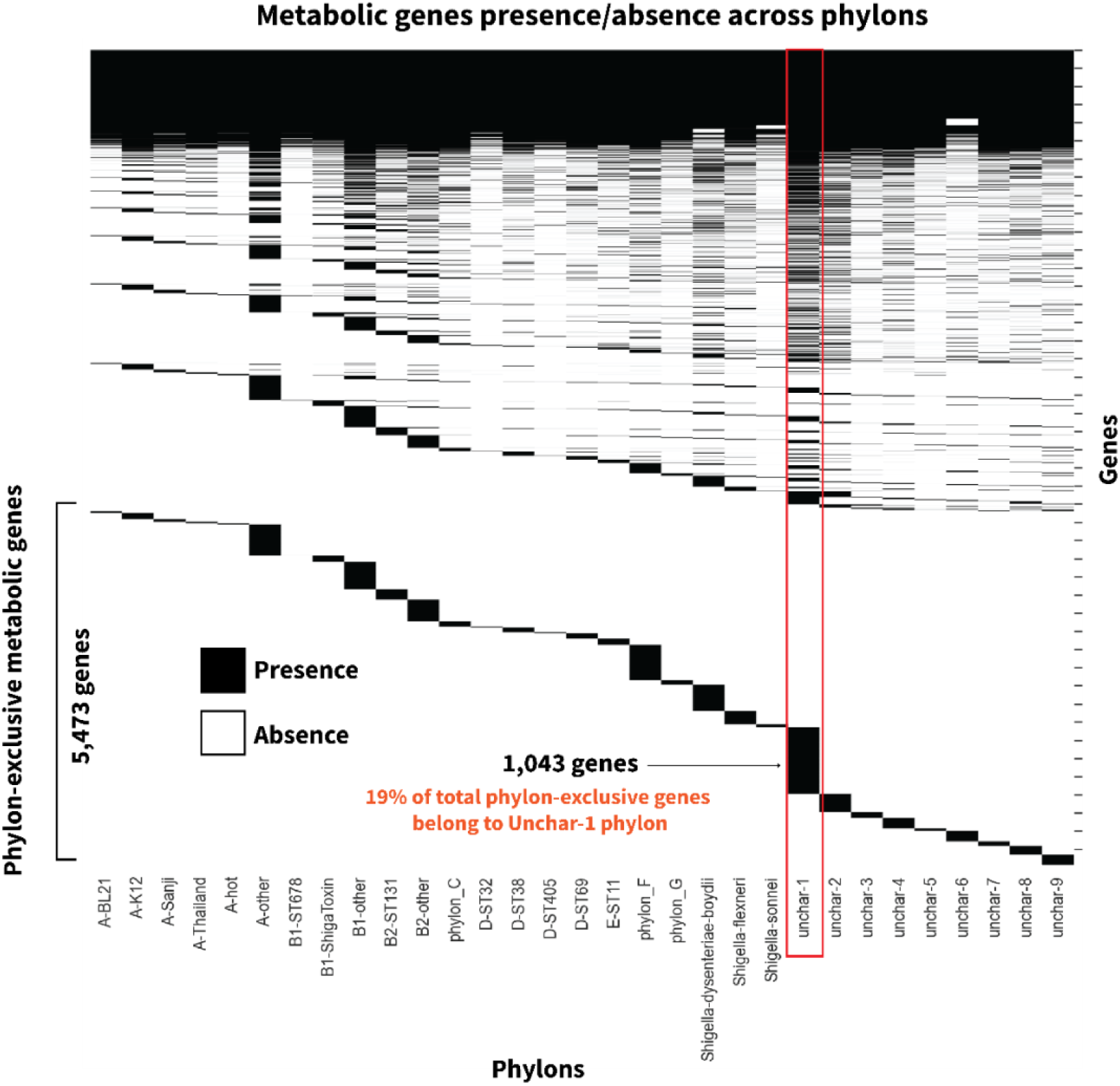
Phylon-based clustering of metabolic genes reveals phylon-exclusive metabolic genes. The heatmap represents the presence or absence of metabolic genes across phylons, with black cells indicating gene presence and white cells indicating gene absence. Genes are displayed on the Y-axis, while phylons are shown on the X-axis. The highlighted phylon, Uncharacterized 1, contains the highest number of phylon-exclusive genes.

**Extended Data Fig 5.**
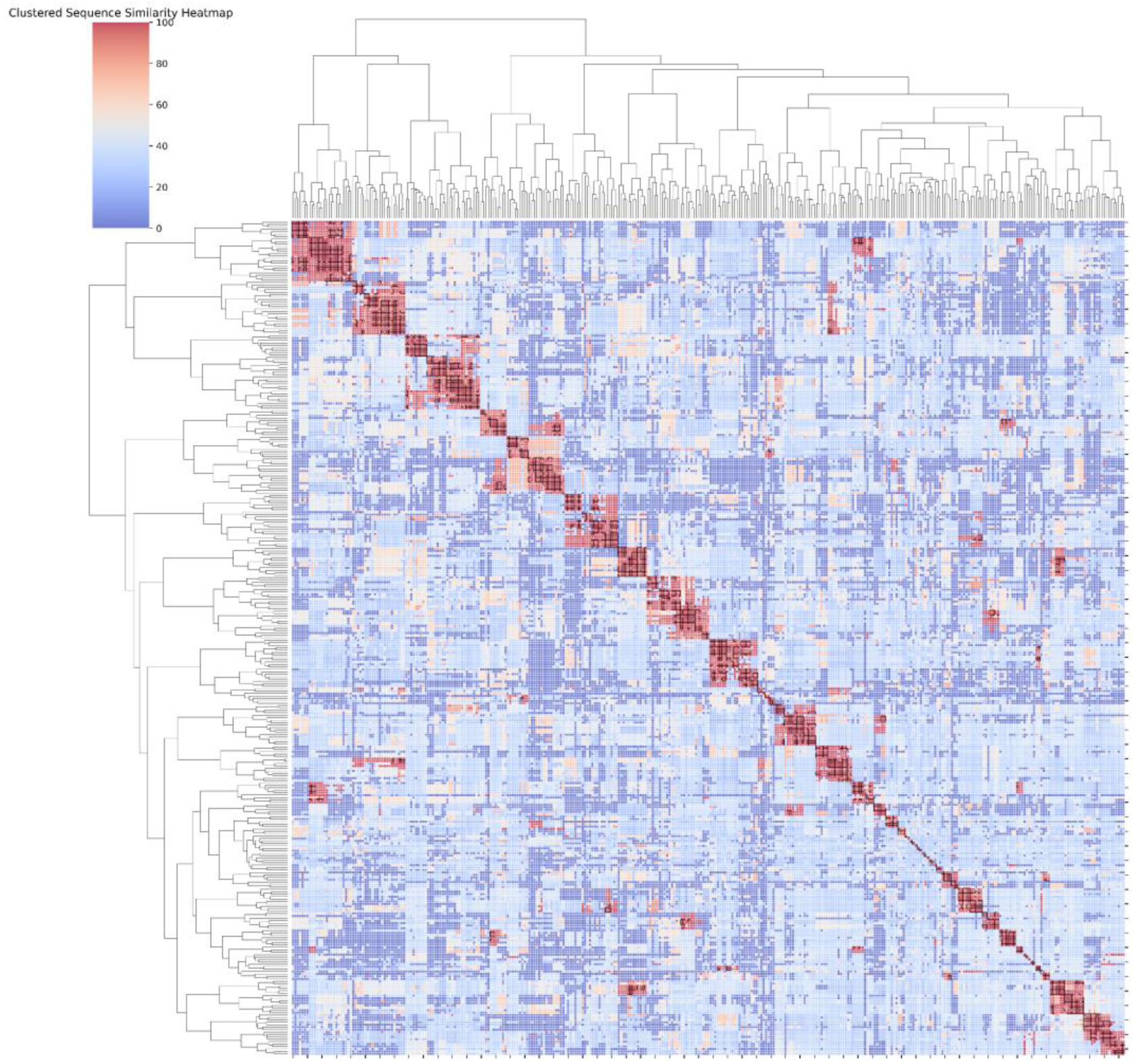
Cyclic di-GMP phosphodiestrase protein sequence similarity. Heatmap represents sequence similarity for Cyclic di-GMP phosphodiestrase genes, for each gene the longest variant was selected to perform a bidirectional BLAST for all pairs of genes, similarity cells are color coded based on their similarity.

**Extended Data Fig 6.**
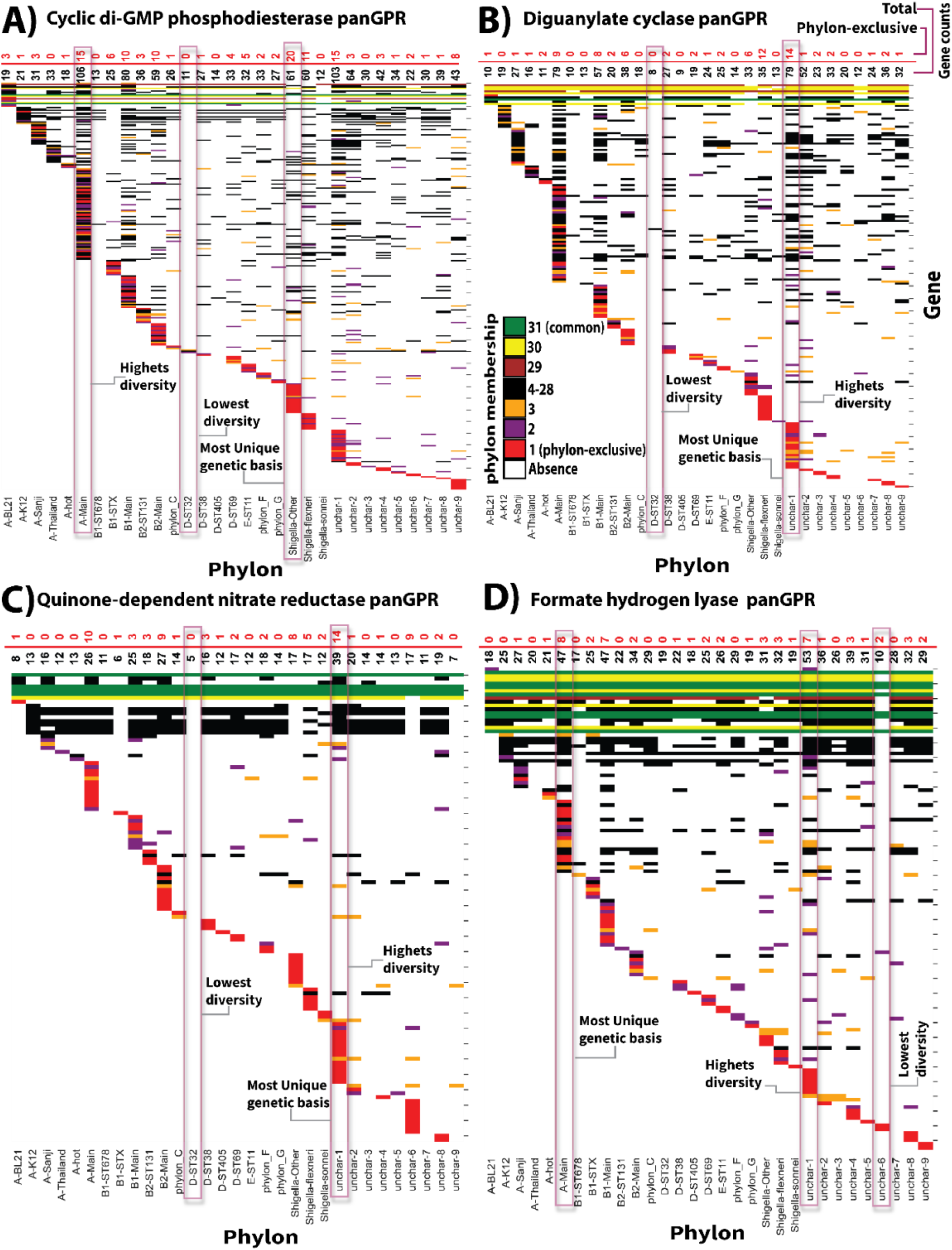
Distribution of Gene Clusters for Reactions with High Genetic Basis Variability Across Phylons. The heatmaps provide a detailed view of the phylon distribution of genes encoding Reactions with High Genetic Basis Variability, color-coded based on the number of phylons in which a gene is found (color coding per inset) for instance red cells show genes that are phylon-exclusive thus cannot be found in other phylons. The annotated texts highlight phylon with highest and lowest variability in genetic basis and also pylons with most unique genetic makeup for each reaction. A) cyclic di-GMP phosphodiesterase (R08991) panGPR. B) Diguanylate cyclase (DGUNC) panGPR. C) Quinon dependent nitrite reductase (NO3R1bpp) panGPR. D) Formate hydrogen lyase (FHL) panGPR.

**Extended Data Fig 7.**
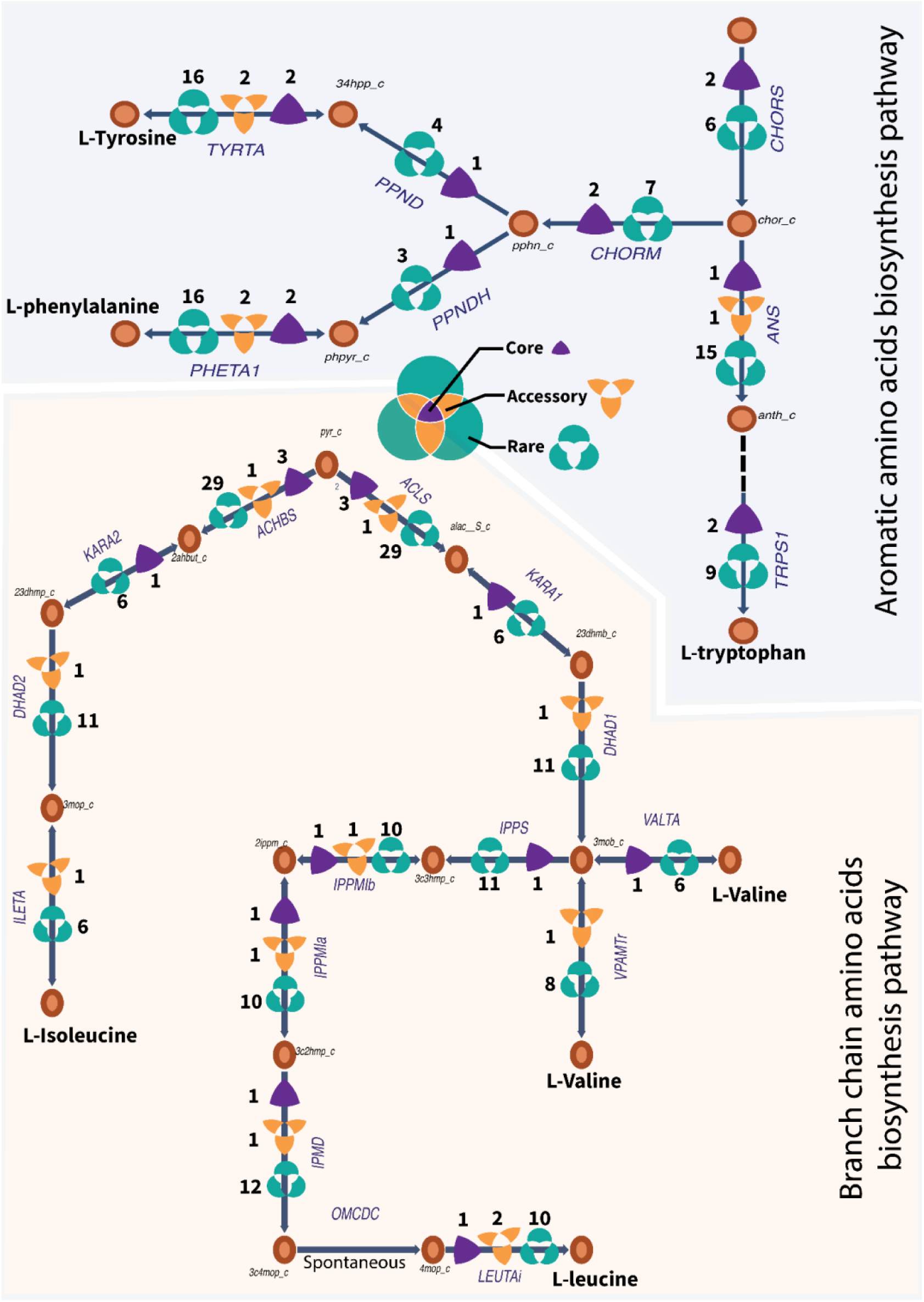
Diversity of genetic basis of reactions in AAA and BCA biosynthesis pathways. Metabolic pathways of AAA biosynthesis and BCA biosynthesis are annotated with the number of core, accessory, and rare genes coding each reaction across the pangenome.

**Extended Data Fig 8.**
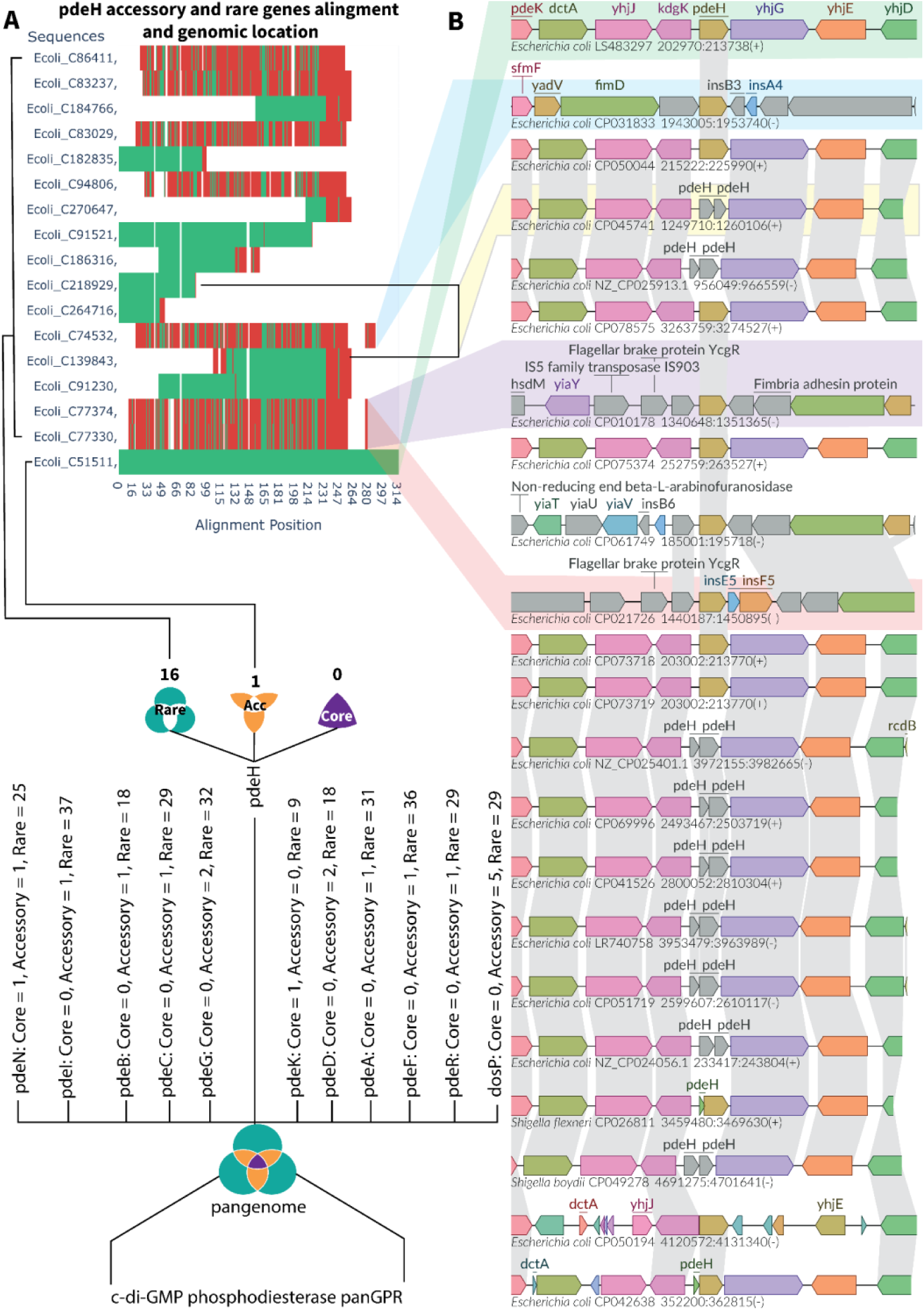
pdeH accessory and rare gene sequence alignment and its genomic neighborhood. **A)** overall view of genes encoding for cyclic di-GMP phosphodiesterase enzyme and sequence alignment for pdeH rare and accessory genes. **B)** genomic neighborhood of accessory and rare pdeH genes, homologous genes are highlighted across genomes. This figure illustrates gene families encoding cyclic di-GMP phosphodiesterase reactions, showing the number of core, accessory, and rare genes within each gene family. The pdeH rare genes exhibit the highest sequence divergence from their reference sequence, which is an accessory gene selected as the reference due to its frequency across strains (in the absence of a core gene). In other gene families, rare genes are primarily fragments of the reference sequence. The pdeN and pdeG families also contain rare genes with substantial differences from their reference (core or accessory) genes. In this analysis, pdeH was selected for an in-depth sequence alignment comparison between rare and reference genes, and to demonstrate the divergent genomic locations of rare genes relative to the reference. In Panel B, the top row shows the conserved genomic location of the reference gene across strains, while the genomic neighborhoods of rare genes vary across strains, often introducing new neighboring genes. These newly introduced genes are annotated upon their first appearance, with each subsequent row indicating entirely new neighboring genes in comparison to the reference gene’s location.

**Extended Data Fig 9.**
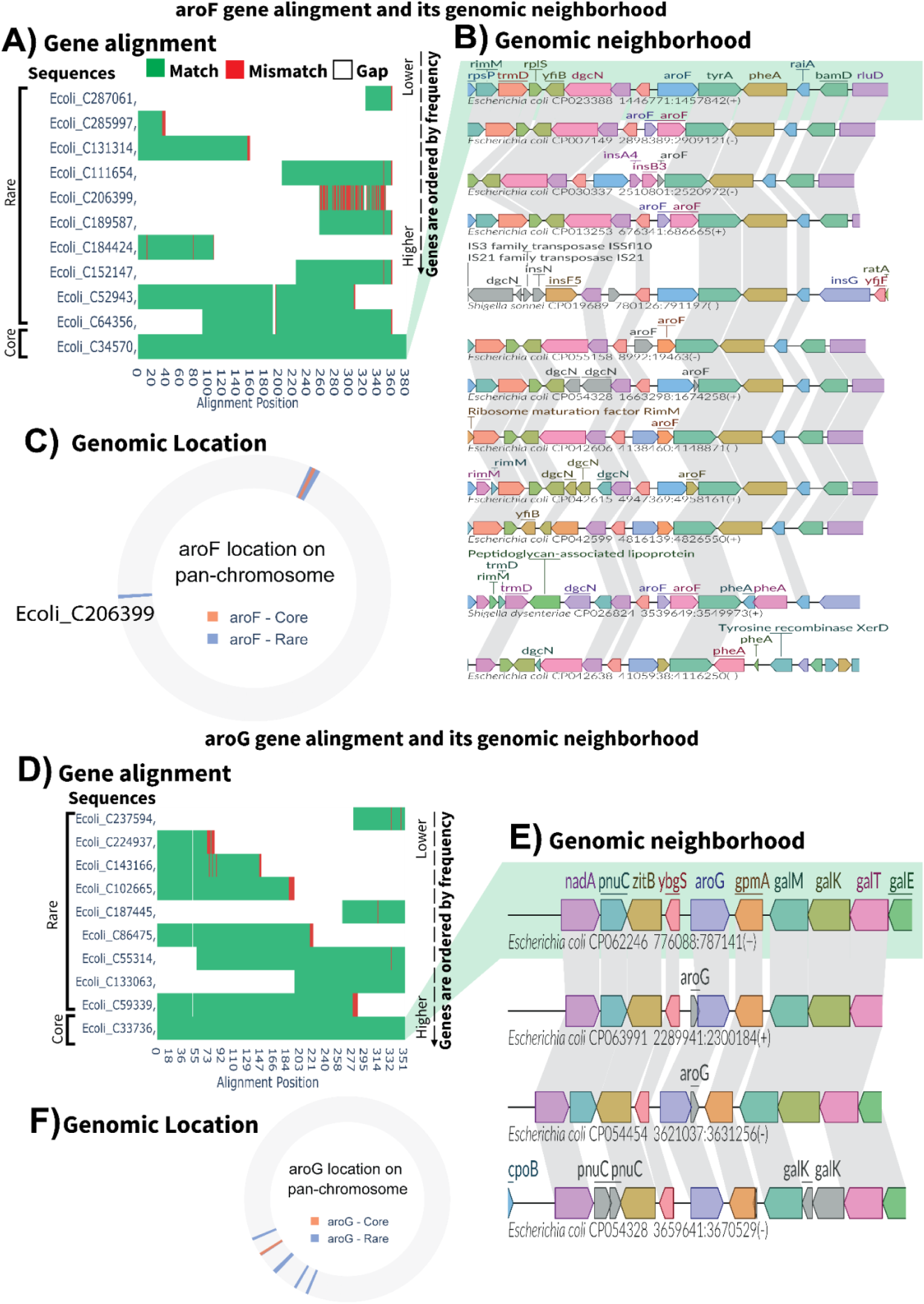
Multiple sequence alignment and gene neighborhood for core, and rare genes of aroG and aroF isozymes. A) MSA of rare aroF genes to core aroH gene, genes are ordered based on frequency from bottom to top. B) genomic neighborhood of core and rare aroF genes, homologous genes are highlighted across genomes. C) Gene locations on pan-chromosome. Core genes are highlighted in orange and Rare genes are in blue. Genomes are aligned by the genes shared by all the genomes and only appear once. D) MSA of rare aroG genes to core aroG gene, genes are ordered based on frequency from bottom to top. E) genomic neighborhood of core and rare aroG genes, homologous genes are highlighted across genomes. F) core genes are highlighted in orange and Rare genes are in blue. Genomes are aligned by the genes shared by all the genomes and only appear once.

**Extended Data Fig 10.**
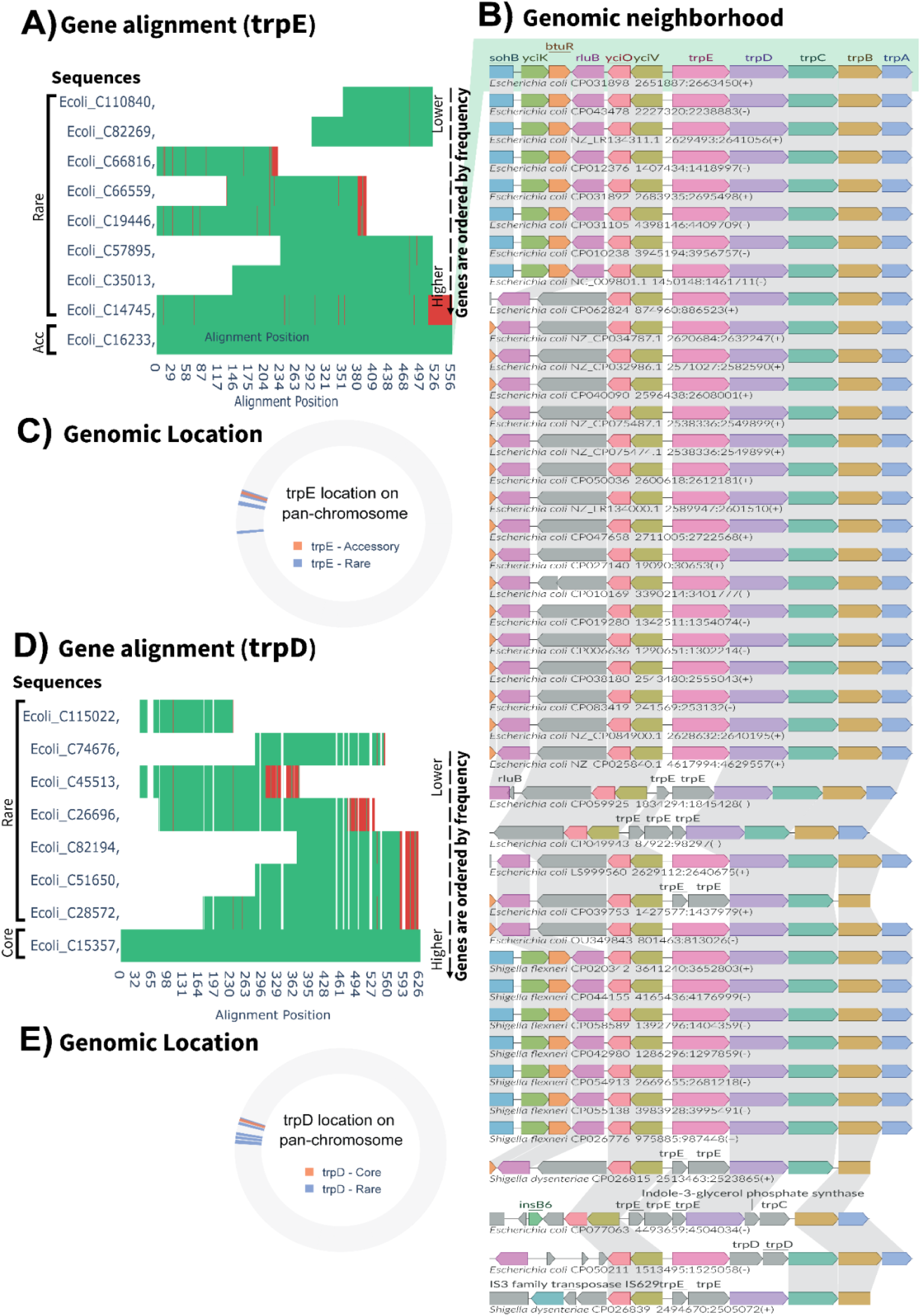
Multiple sequence alignment and gene neighborhood for core, accessory, and rare genes of trpD and trpE isozymes. A) MSA of rare trpE genes to Accessory trpE gene, genes are ordered based on frequency from bottom to top. B) genomic neighborhood of accessory and rare trpE genes and core and rare trpD genes, homologous genes are highlighted across genomes. C) Gene locations on pan-chromosome. Accessory genes are highlighted in orange and Rare genes are in blue. Genomes are aligned by the genes shared by all the genomes and only appear once. D) MSA of rare trpD genes to core trpD gene, genes are ordered based on frequency from bottom to top. E) core genes are highlighted in orange and Rare genes are in blue. Genomes are aligned by the genes shared by all the genomes and only appear once.

**Table S1.** Reaction frequency categories for each phylogroup. Core, accessory, and rare reactions were identified within each phylogroup by grouping GEMs according to their phylogroups. Reaction frequencies were calculated based on the members of each phylogroup, with thresholds of 6.8% and 96.7% applied to classify reactions as rare, accessory, or core, respectively. These categories serve as indicators of the metabolic conservation within each phylogroup.

| Phylogroup | Core | Accessory | Rare |
| --- | --- | --- | --- |
| A | 2465 | 286 | 2 |
| B1 | 2627 | 103 | 23 |
| B2 | 2587 | 164 | 2 |
| C | 2642 | 88 | 23 |
| D | 2582 | 151 | 20 |
| E | 2508 | 189 | 56 |
| E or cladeI | 2753 | 0 | 0 |
| F | 2578 | 133 | 42 |
| G | 2399 | 354 | 0 |
| Shigella | 2295 | 414 | 44 |
| cladeI | 2678 | 32 | 43 |

## Notes

https://zenodo.org/records/13825392

https://github.com/omidard/EcopanGEM

### Competing Interest Statement

The authors have declared no competing interest.

