## Supplementary Table1 for "A Diverse Genetic Basis for Metabolic Reactions is Revealed Through Pangenome analysis"

### **Supplementary Table**

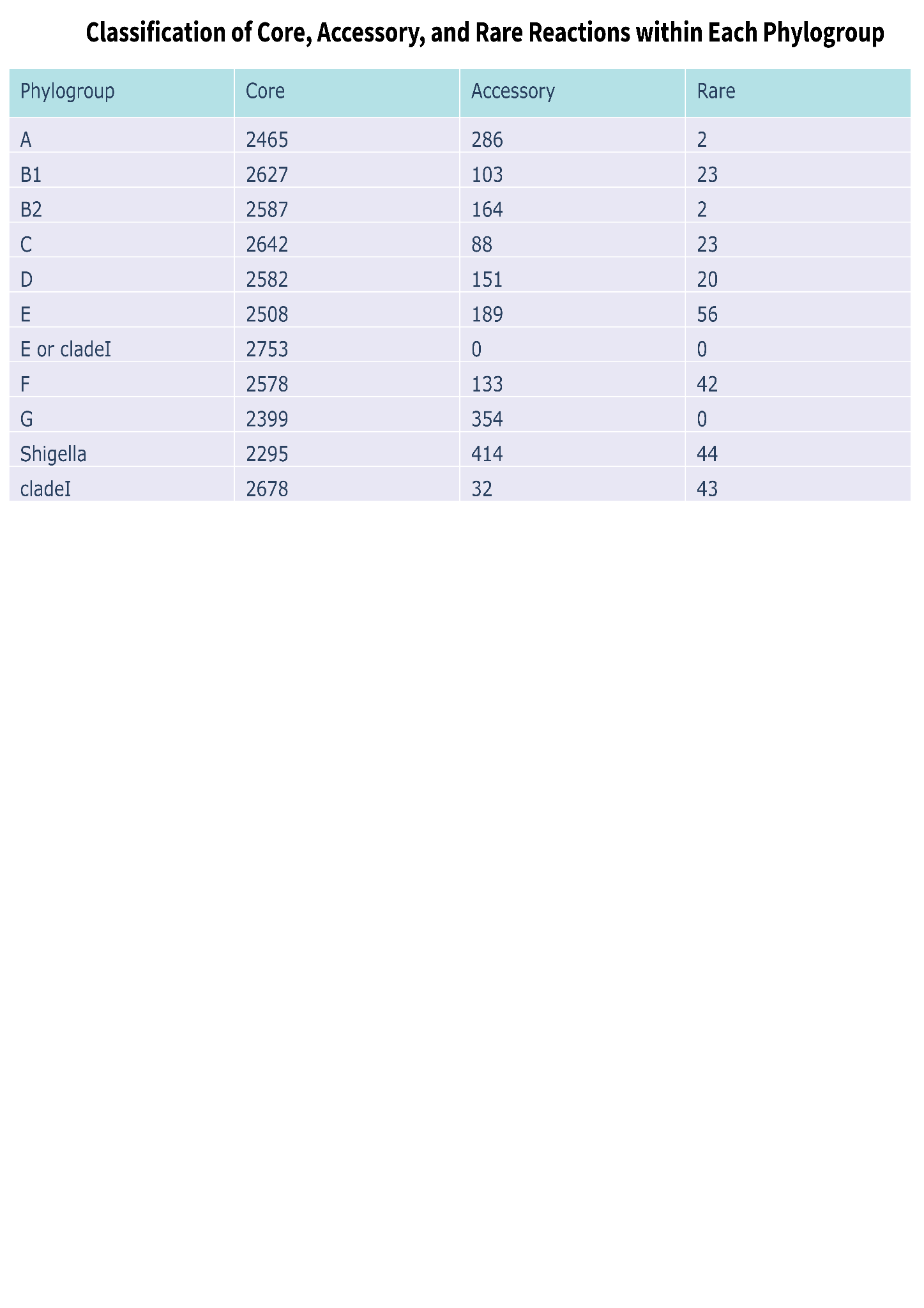

***Table S1. Reaction frequency categories for each phylogroup.*** *Core, accessory, and rare reactions were identified within each phylogroup by grouping GEMs according to their phylogroups. Reaction frequencies were calculated based on the members of each phylogroup, with thresholds of 6.8% and 96.7% applied to classify reactions as rare, accessory, or core, respectively. These categories serve as indicators of the metabolic conservation within each phylogroup.*
